# System-Wide Proteomic Remodeling in Spinal Muscular Atrophy Reveals Tissue-Specific Responses and Partial Rescue by SMN Restoration

**DOI:** 10.64898/2026.03.30.715402

**Authors:** Sofia Vrettou, Stefan Müller, Brunhilde Wirth

## Abstract

Spinal muscular atrophy (SMA), traditionally defined as a neuromuscular disorder characterized by degeneration of lower motor neurons, is increasingly recognized as a multi-organ disease. SMA is caused by deficiency of the survival motor neuron (SMN) protein below a critical threshold required for cellular homeostasis. While motor neurons are particularly vulnerable, the ubiquitous expression and fundamental functions of SMN result in widespread perturbations across multiple tissues.

Here, we generated a label-free quantitative proteomics atlas of spinal cord, heart, and gastrocnemius muscle from wild-type, heterozygous, and SMA mice at the symptomatic stage, including cohorts treated, at postnatal day 1 (P1), with a systemic suboptimal dose of SMN antisense oligonucleotides (SMN-ASOs), resulting in partial SMN restoration. SMN deficiency induced pronounced, tissue-specific proteome remodeling, with peripheral tissues exhibiting broader molecular alterations than spinal cord. Cross-tissue analyses revealed limited overlap, although heart and muscle showed partial convergence in metabolic and mitochondrial-associated pathways. SMN-ASO treatment partially repositioned these proteomes toward control states; however, restoration was incomplete and strongly tissue-dependent, with persistent dysregulation of mitochondrial and metabolic pathways.

These findings demonstrate that SMN deficiency drives systemic yet heterogeneous proteome remodeling and that partial SMN restoration does not fully reverse established molecular alterations.

**Graphical Abstract:** 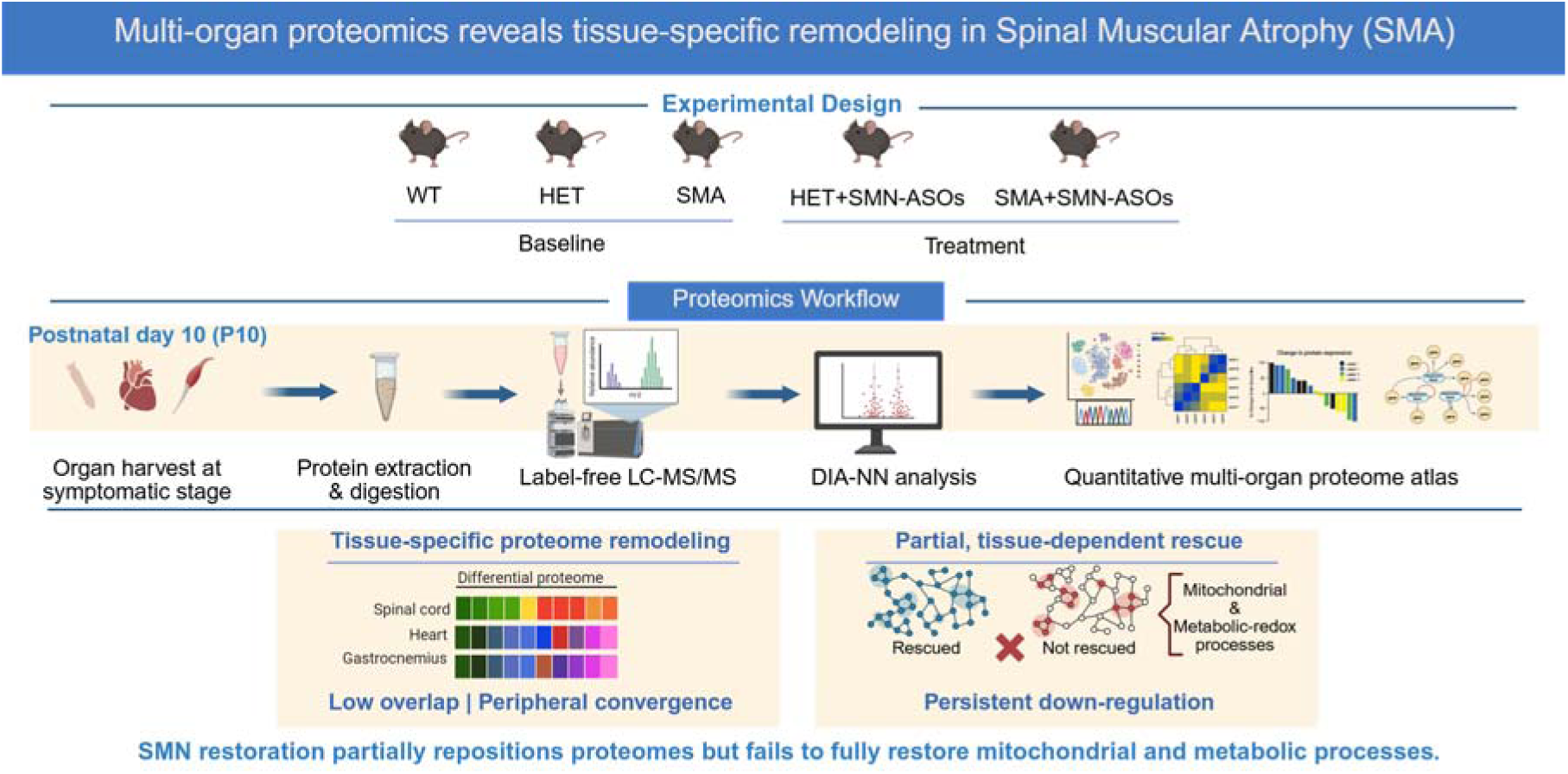

## Introduction

Spinal muscular atrophy (SMA) is caused by deficiency of the survival motor neuron (SMN) protein and is classically defined by progressive loss of lower alpha motor neurons [1–3]. Yet SMN is expressed in virtually all tissues [4], and increasing evidence shows that its deficiency perturbs cellular homeostasis far beyond the nervous system [5, 6]. Peripheral organs, including skeletal muscle and heart, exhibit early and sustained abnormalities, underscoring that SMA is not confined to the motor unit but represents a systemic disorder with tissue-specific manifestations [7–11].

A central challenge in SMA research is understanding why organs respond so differently to a shared molecular deficit. Neuromuscular tissues are particularly vulnerable, but even among them, the magnitude and nature of molecular remodeling vary markedly [6, 12, 13]. Spinal cord pathology dominates clinical presentation, while skeletal and cardiac muscles show intrinsic metabolic and mitochondrial alterations that cannot be explained by denervation alone [8, 14–18]. These observations suggest that SMN deficiency engages organ-specific adaptive and maladaptive programs shaped by developmental timing, metabolic demand, and compensatory capacity. Whether these divergent responses are reflected in coordinated, tissue-specific proteomic programs remain unresolved.

The advent of SMN-restoring therapies, including antisense oligonucleotides (ASOs), has fundamentally improved the clinical course of SMA, particularly when treatment is initiated early [19–23]. However, therapeutic benefit is not uniform, and molecular normalization across tissues is often incomplete [24]. This raises an important question: whether early SMN restoration normalizes tissue proteomes or instead establishes distinct, partially corrected states remains unclear.

Unbiased proteomics offers a powerful means to address this question by capturing coordinated protein-level changes across pathways and cellular compartments. While recent targeted and single-organ studies have demonstrated organ-dependent molecular remodeling and partial correction following SMN-ASO treatment [24–26], a unified, cross-tissue proteomic analysis spanning both neuronal and muscular compartments within the same experimental framework is lacking.

Here, we present a label-free quantitative proteomics atlas of spinal cord, gastrocnemius muscle, and heart from wild-type (WT), heterozygous (HET), and SMA mice at the symptomatic stage (postnatal day 10; P10), including cohorts treated with SMN-ASOs at postnatal day 1 (P1), which led in partial SMN restoration. By jointly examining central and peripheral neuronal and muscular tissues within a unified experimental design, we define organ-specific proteomic signatures of SMN deficiency and evaluate how early SMN restoration reshapes these molecular states. This atlas establishes a cross-tissue proteomic framework for interpreting systemic SMN biology and for guiding future mechanistic and therapeutic studies in SMA.

## Materials and Methods

### Animal Model and Tissue Collection

Severe Taiwanese SMA mice (Smn^−/−^; SMN2^tg/0^), heterozygous carriers (Smn^+/−^; SMN2^tg/0^), and wild-type (WT) controls were generated and maintained as previously described [24, 26–29].

For SMN restoration, neonatal mice received a single subcutaneous injection of a splice-modulating SMN-targeting antisense oligonucleotide (SMN-ASO) at P1, as previously described based on validated protocols [24–26, 28, 30, 31]. This oligonucleotide acts on *SMN2* pre-mRNA splicing to favor exon 7 inclusion and increase expression of full-length SMN protein [32]. Because the SMN-ASO was administered systemically at a suboptimal dose, the resulting rescue was partial and not expected to fully normalize SMN levels across tissues [24, 33]. Control WT animals were not injected. Treated and untreated animals were sacrificed at P10, and tissues were rapidly dissected, snap-frozen in dry ice, and stored at −80 °C until processing.

Mice were housed under controlled environmental conditions with a 12-hour light/dark cycle and had unrestricted access to food and water. All breeding, housing, and experimental procedures were carried out in a specific pathogen-free environment. Animal experiments were conducted in compliance with applicable institutional and governmental guidelines and were approved by the Landesamt für Natur, Umwelt und Verbraucherschutz Nordrhein-Westfalen (LANUV) under application numbers: 81-02.04.2020.A196, 81-02.04.2019.A017, 81-02.04.2019.A138, §4.23.008, and §4.22.002.

### Protein Extraction and Digestion

Protein extraction, reduction, alkylation, and enzymatic digestion were performed using a standardized urea-based workflow as previously described [25, 26]. Briefly, tissues were homogenized in 8 M urea buffer (50 mM TEAB, pH 8.0) supplemented with protease inhibitors. Protein lysates were reduced, alkylated, and sequentially digested with Lys-C and trypsin. Peptides were acidified and desalted using SDB-RPS StageTips prior to LC–MS/MS analysis.

### LC–MS/MS Data Acquisition and DIA-NN Processing

Peptides were analyzed by nano LC–MS/MS on a Vanquish Neo system coupled to a Thermo Orbitrap Exploris 480 mass spectrometer equipped with a FAIMS Pro interface, as previously described [26]. Data-independent acquisition (DIA) was performed across the mass range of 400-1000 m/z. Instrument parameters and acquisition settings were kept consistent across all tissues to ensure comparability.

Raw DIA data were processed using DIA-NN (v1.8.1) [34] with a predicted spectral library generated from the Mus musculus UniProt canonical database, as described previously [26]. Data were filtered at 1% false discovery rate (FDR) at both precursor and protein levels. Protein-level label-free quantification (LFQ) intensities were exported for downstream analysis.

Comprehensive instrument parameters and DIA-NN processing settings are described in detail in our accompanying Data in Brief publication on liver proteomics from the same animal model, ensuring methodological transparency and reproducibility [26].

### Bioinformatics Analysis

#### 1. Statistical Analysis

Statistical analysis was performed in Perseus (v1.6.15) [35] following established workflows [25, 26]. Protein intensities were log2-transformed, filtered for valid values across biological replicates, and missing values were imputed. Differential expression was assessed using two-sample t-tests with permutation-based FDR correction (FDR = 0.05; S0 = 0.1 unless otherwise specified). Principal component analysis was performed on processed datasets. The number of biological replicates per group is indicated in the corresponding figures and legends.

#### 2. Proteomic Stratification and Cross-Organ Integration Analysis

Because proteomic responses to SMN-ASO treatment were heterogeneous within the SMA injected with SMN-ASOs (SMA+ASO) group, treated SMA samples were stratified according to the degree of proteome repositioning observed by principal component analysis (PCA) in each tissue. Samples showing a more pronounced shift away from untreated SMA and toward the WT/HET proteomic space were considered to exhibit greater proteomic normalization, whereas samples remaining closer to the untreated SMA cluster were considered to exhibit limited proteomic normalization. By contrast, HET+ASO samples showed comparatively homogeneous proteomic profiles across animals within each tissue and were therefore analyzed as a single treatment group without additional stratification.

Using this stratified framework, we next assessed the extent to which SMN-ASO treatment directionally normalized proteins significantly altered in SMA relative to WT. First, we sorted the proteins identified as significantly dysregulated in the WT versus SMA comparison and then evaluated their convergence (presence or absence; fold change; significance thresholds) in the SMA versus SMA+ASO comparison. Proteins were considered directionally rescued when their abundance shifted toward WT levels after SMN-ASO treatment. Proteins that remained significantly altered without directional reversal were classified as persistent. Proteins significantly dysregulated in WT versus ASO but not detected within the cohort of significantly altered in SMA versus SMA+ASO were classified as no response in ASO. Proteins significantly altered in the SMA versus SMA+ASO comparison but not identified as significantly dysregulated in the WT versus SMA comparison were classified as ASO-responsive only.

Cross-organ integration was performed using UniProt identifiers to determine shared and tissue-specific protein alterations. Analyses focused on proportional rescue and pathway convergence across tissues.

#### 3. Network and Functional Enrichment Analysis

Functional enrichment analysis was performed using STRING (v11.5; Mus musculus) and visualized in Cytoscape with the ClueGO plugin. Gene Ontology (Biological Process, Molecular Function, Cellular Component), KEGG, and Reactome databases were used for pathway analysis. Statistical significance was determined using two-sided hypergeometric testing with Benjamini-Hochberg correction (adjusted p < 0.05). A kappa score threshold of 0.4 was applied for functional grouping.

#### 4. Data Visualization

Volcano plots were generated using InstantClue [36] based on statistical outputs from Perseus. PCA and rescue classification visualizations were generated using SRplot [37]. Visualization tools were used exclusively for graphical representation and not for statistical analysis.

## Results

### 1. Global proteome profiling reveals organ-specific remodeling in SMA

To define baseline proteome alterations associated with SMN deficiency across neuronal and muscular tissues, label-free quantitative proteomics was performed on spinal cord, heart, and gastrocnemius muscle isolated at P10 from WT, HET, and SMA mice. The goal was to quantitatively compare the extent and nature of proteome remodeling across neuromuscular tissues. After quality control and exclusion of samples with insufficient signal or genotype-inconsistent clustering, final cohort sizes were: spinal cord (4 WT, 5 HET, 4 SMA), heart (4 WT, 5 HET, 5 SMA), and gastrocnemius (5 WT, 5 HET, 4 SMA).

Across tissues, 4,983 proteins were identified in spinal cord, 2,912 in heart, and 3,248 in gastrocnemius. After filtering for valid quantification, 4,787 proteins were retained in spinal cord, 2,761 in heart, and 3,164 in gastrocnemius.

PCA revealed genotype-dependent separation across all three tissues (Figure 1A-C). In spinal cord and gastrocnemius, SMA samples segregated from WT, with HET samples positioned intermediately, consistent with SMN dosage-dependent proteome remodeling (Figure 1A, C). In heart, clear separation between genotypes was observed (Figure 1B). Unsupervised hierarchical clustering of global protein abundance (Figure 1D-F) supported these findings, as samples grouped primarily according to genotype across tissues.

**Figure 1.**
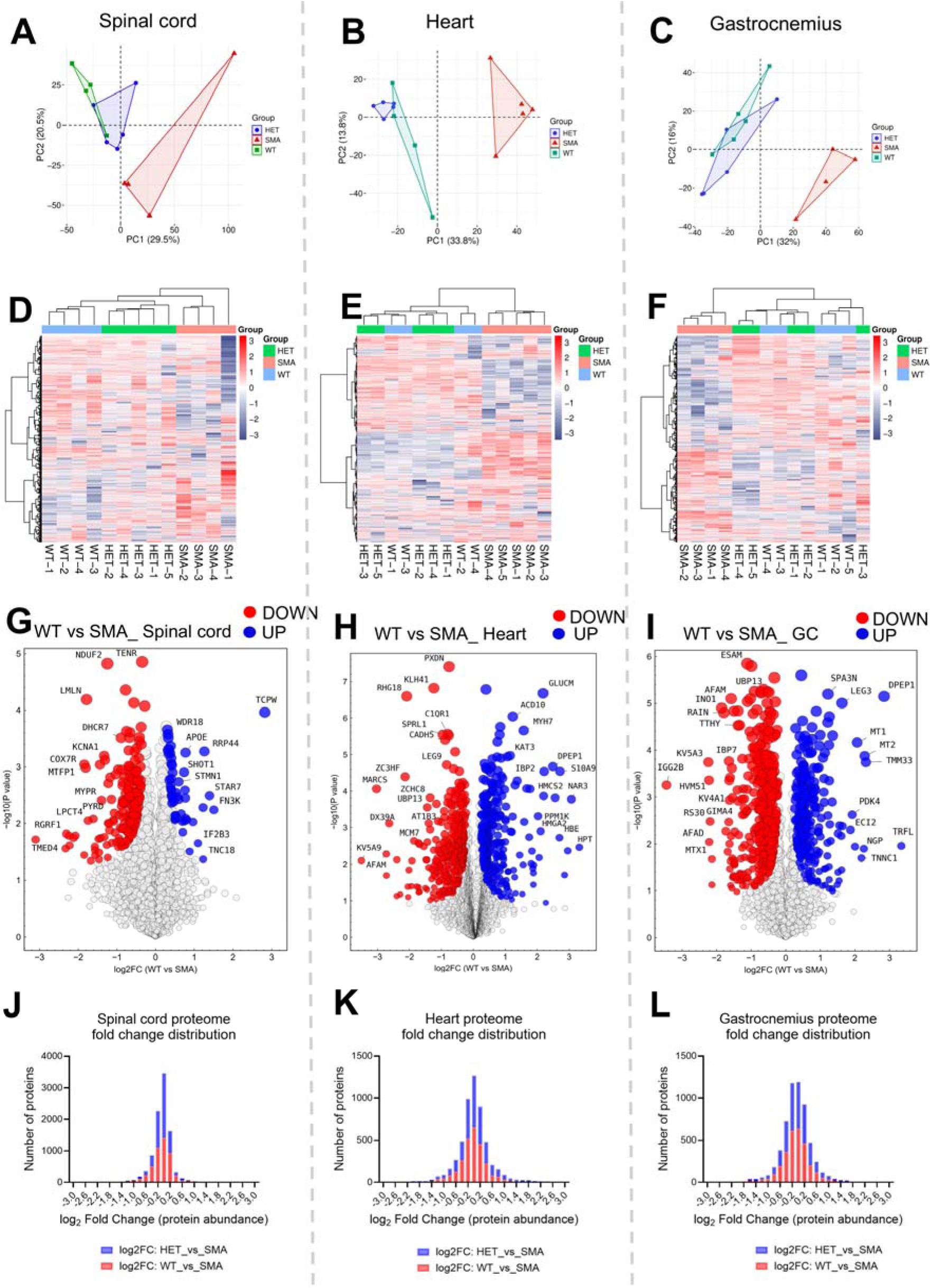
Global proteome remodeling across neuromuscular tissues in SMA. (A-C) Principal component analysis (PCA) of label-free quantified proteomes from spinal cord (A), heart (B), and gastrocnemius muscle (C) at P10, showing genotype-dependent separation among WT, HET, and SMA samples. (D-F) Unsupervised hierarchical clustering of protein abundance across the same tissues, demonstrating sample grouping primarily by genotype. (G-I) Volcano plots of differential protein abundance (WT vs SMA) generated in Perseus (FDR = 0.05, S0 = 0.1), highlighting significantly downregulated (red) and upregulated (blue) proteins in spinal cord (G), heart (H), and gastrocnemius (I). (J-L) Distribution of log2 fold changes in protein abundance, illustrating the magnitude and direction of proteome-wide alterations, with broader shifts observed in heart and gastrocnemius relative to spinal cord. Complete statistical outputs are provided in Supplementary Tables S1–S5.

Differential abundance analysis between WT and SMA was performed using Perseus (FDR 0.05, S0 = 0.1). Volcano plots (Figure 1G-I) revealed marked differences in the extent of proteome remodeling across tissues. Spinal cord exhibited comparatively limited proteome remodeling (Figure 1G), whereas heart and gastrocnemius displayed substantially broader sets of differentially abundant proteins (Figure 1H-I). Fold-change distribution plots (Figure 1J-L) summarize the global magnitude and direction of genotype-dependent shifts, demonstrating broader distribution changes in heart and gastrocnemius compared to spinal cord. Complete lists of significantly altered proteins for all comparisons are provided in the Supplementary Tables S1-S3.

Volcano plots for HET versus SMA are shown in Supplementary Figure 1. In spinal cord, no proteins passed the applied Perseus thresholds (Supplementary Figure S1A), consistent with the PCA positioning of HET samples in this tissue (Figure 1A). In contrast, heart and gastrocnemius exhibited significant alterations between HET and SMA (Supplementary Figure S1B-C). Comparison of WT versus HET revealed modest differences overall, with limited significant proteins detected (Supplementary Figure S1G-H).

Intersection analysis across tissues demonstrated that proteome alterations were predominantly tissue-specific (Supplementary Figure S1G-H; Supplementary Tables S4-S5), with only limited overlap between spinal cord and peripheral tissues. By contrast, heart and gastrocnemius showed partial convergence in membrane-organization, metabolic and mitochondrial-associated pathways (Supplementary Figure S2A).

To define the cellular architecture affected by SMN deficiency, 1D enrichment analysis was performed for WT versus SMA in each tissue (Figure 2A-C). In spinal cord, enriched categories included membrane-associated, nuclear, endoplasmic reticulum, and mitochondrial components (Figure 2A), supporting increasing recognition of mitochondrial involvement in SMA pathology [17, 18]. In heart, enrichment involved mitochondrial structures, ribosomal compartments, extracellular matrix–associated elements, and cytoskeletal assemblies (Figure 2B), consistent with the diverse molecular roles attributed to SMN in RNA metabolism [4, 38–40] and cytoskeletal regulation [41–45]. Likewise, in gastrocnemius, cytosolic, ribosomal, mitochondrial, and contractile-associated components were enriched (Figure 2C).

**Figure 2.**
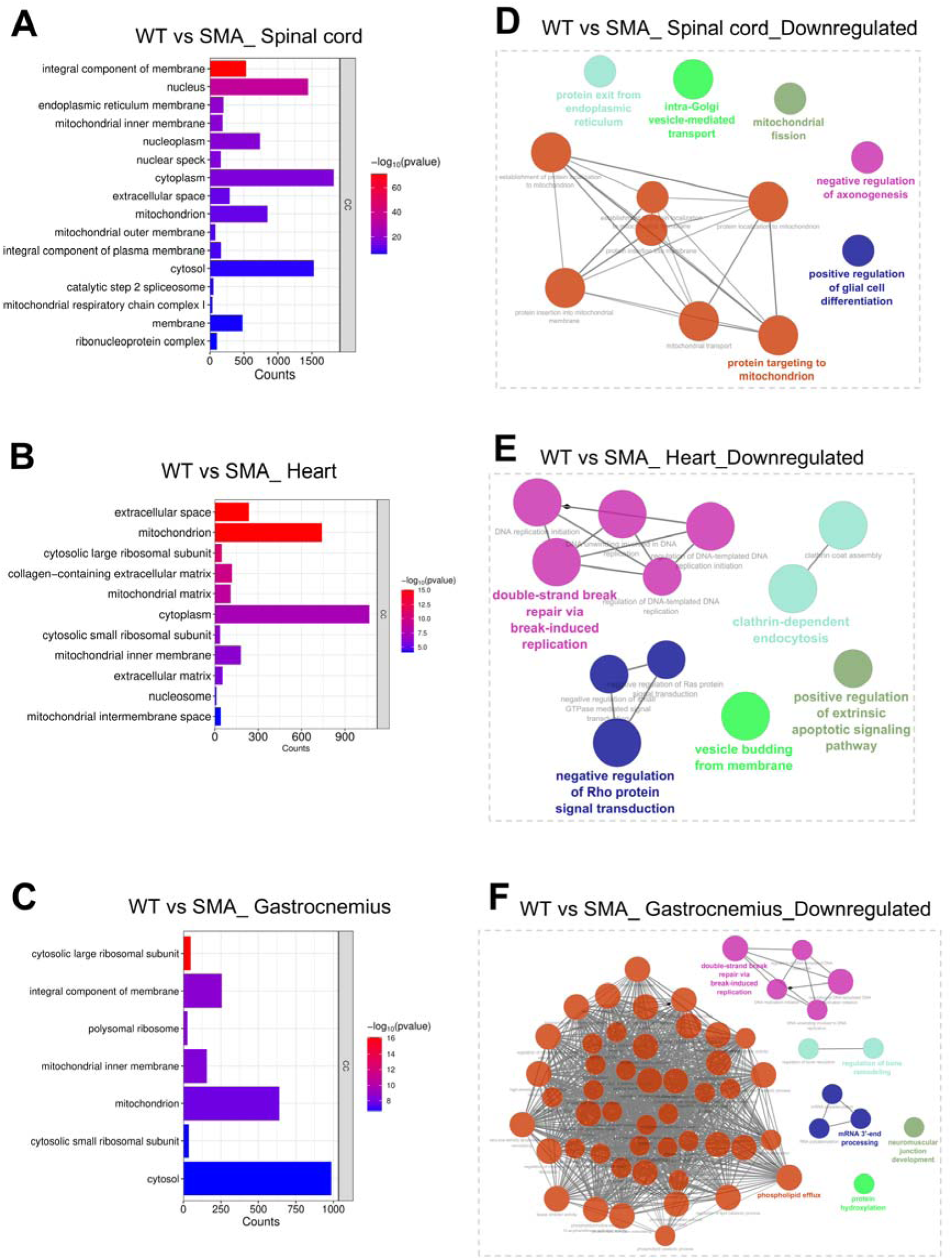
Functional enrichment and network organization of downregulated proteomes in WT versus SMA across tissues. (A-C) One-dimensional enrichment analysis (Perseus) of cellular component categories (GOCC) for significantly downregulated proteins (WT vs SMA) in spinal cord (A), heart (B), and gastrocnemius muscle (C), highlighting subcellular compartments affected by SMN deficiency. (D-F) STRING network analysis of significantly downregulated proteins in spinal cord (D), heart (E), and gastrocnemius (F), visualized in Cytoscape with ClueGO functional grouping. Networks reveal tissue-specific clustering of biological processes, including mitochondrial, endoplasmic reticulum and Golgi-associated pathways, axonogenesis and glial differentiation in spinal cord (D); DNA repair, intracellular trafficking, and clathrin-mediated endocytosis in heart (E); and phospholipid metabolism, mRNA processing, DNA repair, and neuromuscular junction-related pathways in gastrocnemius (F).

STRING network analysis of significantly downregulated proteins in WT versus SMA comparisons (Figure 2D-F) revealed tissue-specific organization. In spinal cord, downregulated proteins formed interconnected clusters associated with mitochondrial components, endoplasmic reticulum and Golgi compartments, axonogenesis, and glial cell differentiation (Figure 2D). In heart, downregulated proteins clustered in pathways related to DNA repair mechanisms, intracellular trafficking processes as well as clathrin-mediated endocytosis (Figure 2E), consistent with previous evidence linking endocytic dysregulation to SMA pathophysiology [33, 46–49]. In gastrocnemius, downregulated networks included phospholipid efflux pathways, DNA repair processes, mRNA processing, and neuromuscular junction development (Figure 2F), reflecting established neuromuscular junction vulnerability in SMA [50–53]. STRING network analysis of significantly upregulated proteins in WT versus SMA comparisons is shown in Supplementary Figure S3.

A parallel 1D enrichment and STRING analysis was performed for HET versus SMA comparisons (Figure 3; Supplementary Figure S4). In spinal cord, STRING analysis revealed mitochondrial-associated clusters (Figure 3A, D) similar to those observed in WT versus SMA (Figure 2A, D). In heart, enrichment was dominated by extracellular and ribosomal components (Figure 3B), and STRING networks showed prominent spliceosomal and snRNP-associated clusters together with endosome and vesicle localization pathways (Figure 3E). In gastrocnemius, enrichment involved mitochondrial and ribosomal compartments (Figure 3C), and STRING analysis identified a dominant phospholipid-associated cluster (Figure 3F) mirroring the WT versus SMA analysis (Figure 2C, F). Upregulated protein networks for HET versus SMA comparisons are shown in Supplementary Figure S4.

**Figure 3.**
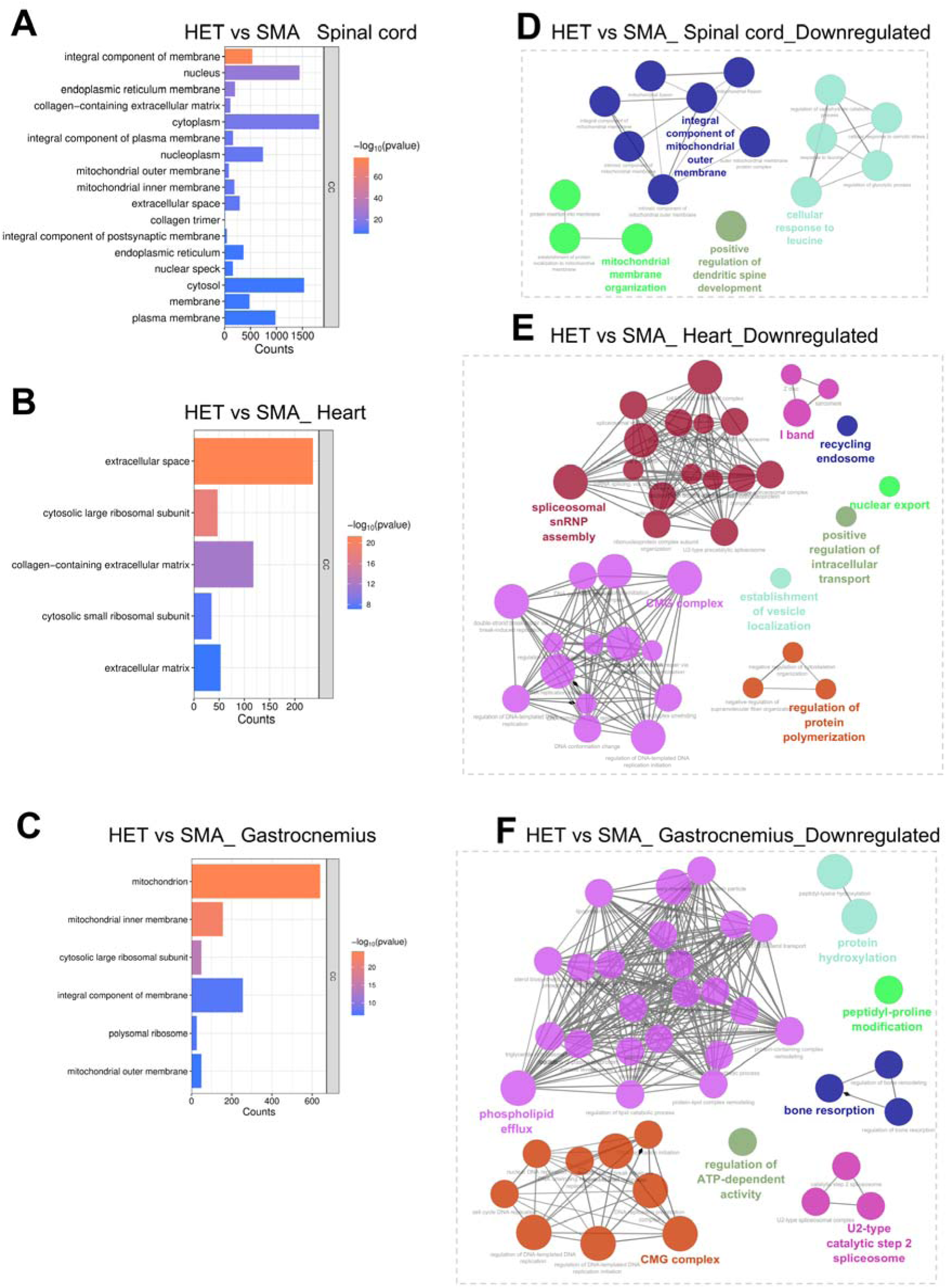
Functional enrichment and network organization of downregulated proteomes in HET versus SMA across tissues. (A-C) One-dimensional enrichment analysis (Perseus) of cellular component categories (GOCC) for significantly downregulated proteins (HET vs SMA) in spinal cord (A), heart (B), and gastrocnemius muscle (C), identifying subcellular compartments associated with SMN dosage-dependent proteome alterations. (D-F) STRING network analysis of significantly downregulated proteins in spinal cord (D), heart (E), and gastrocnemius (F), visualized in Cytoscape with ClueGO functional grouping. Networks reveal tissue-specific clustering of biological processes, including mitochondrial and membrane-associated components in spinal cord (D); spliceosomal, ribonucleoprotein, and vesicle transport pathways in heart (E); and phospholipid metabolism, mitochondrial organization, and RNA-related processes in gastrocnemius (F).

Together, these analyses demonstrate that SMN deficiency induces clear, genotype-dependent proteome remodeling at the symptomatic stage P10. These changes are largely tissue-specific, with partial overlap between heart and gastrocnemius, and involve recurrent mitochondrial, trafficking, and RNA-related pathways.

### 2. Organ-specific pathway responses to SMN-ASO treatment

#### 2.1. Global proteomic repositioning following partial SMN restoration

To assess whether SMN elevation alleviates tissue-specific proteome alterations, we analyzed spinal cord, heart, and gastrocnemius muscle samples from SMN-ASO-treated animals. Because proteomic responses within the SMA+ASO group were heterogeneous, subsequent analyses focused on the SMA+ASO samples showing the clearest proteomic shift away from untreated SMA and toward the WT/HET state in each tissue based on PCA and as it is discussed in Bioinformatics analysis section. This approach was used to define organ-specific molecular responses to partial SMN restoration.

PCA demonstrated organ-dependent responses to SMN-ASO treatment. In spinal cord (Figure 4A), heart (Figure 4B), and gastrocnemius (Figure 4C), SMA+ASO samples exhibited variable degrees of proteomic repositioning relative to untreated SMA, with a subset of samples shifting toward the WT/HET proteomic space in each tissue. Even in that subgroup, repositioning was incomplete across all tissues, indicating partial rather than full proteome normalization following SMN elevation.

**Figure 4.**
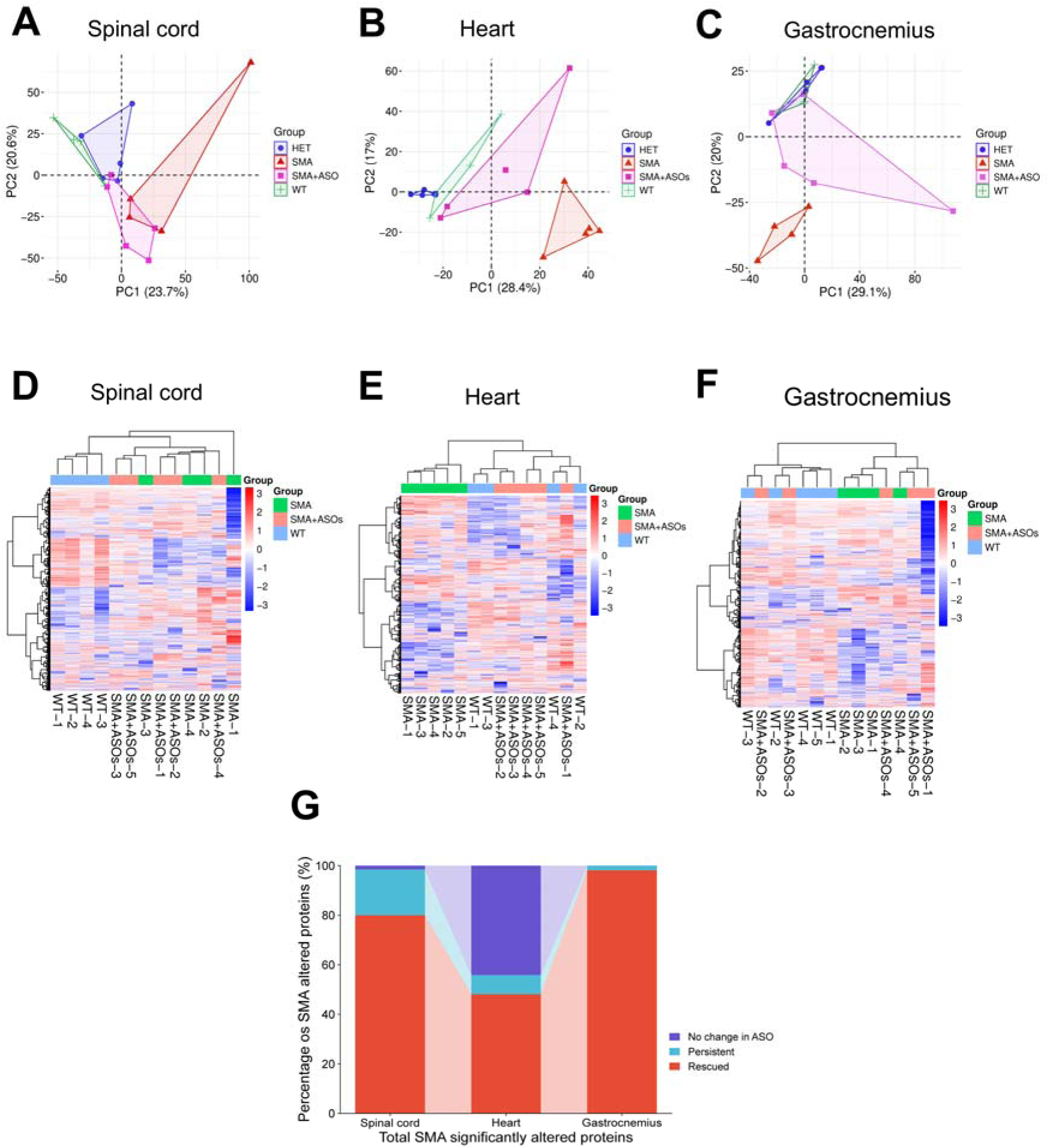
Organ-specific proteome repositioning and protein response classification following SMN-ASO treatment. (A-C) Principal component analysis (PCA) of spinal cord (A), heart (B), and gastrocnemius muscle (C) proteomes including WT, HET, SMA, and SMA+ASO groups, showing organ-dependent repositioning of ASO-treated samples relative to untreated SMA. (D-F) Unsupervised hierarchical clustering of protein abundance across the same tissues, illustrating tissue-specific patterns of partial proteome normalization and persistence following SMN partial restoration. (G) Classification of SMA-associated proteins based on directional response to SMN-ASO treatment, shown as proportions of rescued (shifted toward WT), persistent (unchanged), and no change in ASO across tissues. Detailed target distribution per category and organ is provided in Supplementary Table S6.

Hierarchical clustering and heatmap analysis further illustrated tissue-specific patterns of proteomic normalization and persistence (Figure 4D-F).

To systematically quantify treatment effects, proteins significantly altered in SMA relative to WT were classified according to their response to SMN-ASO treatment as rescued (shifted toward WT levels), persistent (remaining altered without directional reversal), or no change in ASO (when no significant treatment-associated change was detected). The relative distribution of these categories per organ is summarized in Figure 4G. ASO-responsive-only corresponds to significantly altered proteins following treatment but not significantly dysregulated in the WT versus SMA comparison. Complete protein lists for each classification in spinal cord, heart, and gastrocnemius are provided in Supplementary Tables S6-S9.

To contextualize SMA-specific effects, PCA and heatmap analyses including WT, HET, SMA, HET+ASO, and SMA+ASO groups are shown in Supplementary Figure S5A-I. HET samples clustered closely with WT across tissues, and HET+ASO samples did not exhibit major global shifts relative to untreated HET controls (Supplementary Figure S5A-C). These findings indicate that SMN-ASO treatment exerts minimal proteomic effects in non-diseased contexts. Differential expression analysis between HET and HET+ASO groups was performed for each organ. Volcano plots are shown in Supplementary Figure S5G-I. Across spinal cord, heart, and gastrocnemius, a limited number of proteins were significantly altered following ASO administration, with significant changes detected only in spinal cord (Supplementary Figure S5G). The magnitude of change was modest compared to the SMA versus SMA+ASO comparison. Notably, HET+ASO samples displayed comparatively homogeneous clustering behavior across tissues and were therefore considered as a single treatment group without further stratification.

#### 2.2. Functional enrichment of rescued and persistent proteomic responses

To further define the molecular processes responsive or resistant to SMN elevation, pathway enrichment analysis was performed separately for proteins rescued from SMA-associated upregulation and downregulation (Figure 5A-B; Supplementary Figure S6A). Because the ASO-responsive-only category reflects treatment-associated changes outside the set of baseline SMA-dysregulated proteins, downstream functional enrichment in the main analysis focused on rescued and persistent categories, while ASO-responsive-only proteins are provided in Supplementary Tables S6-S9.

**Figure 5.**
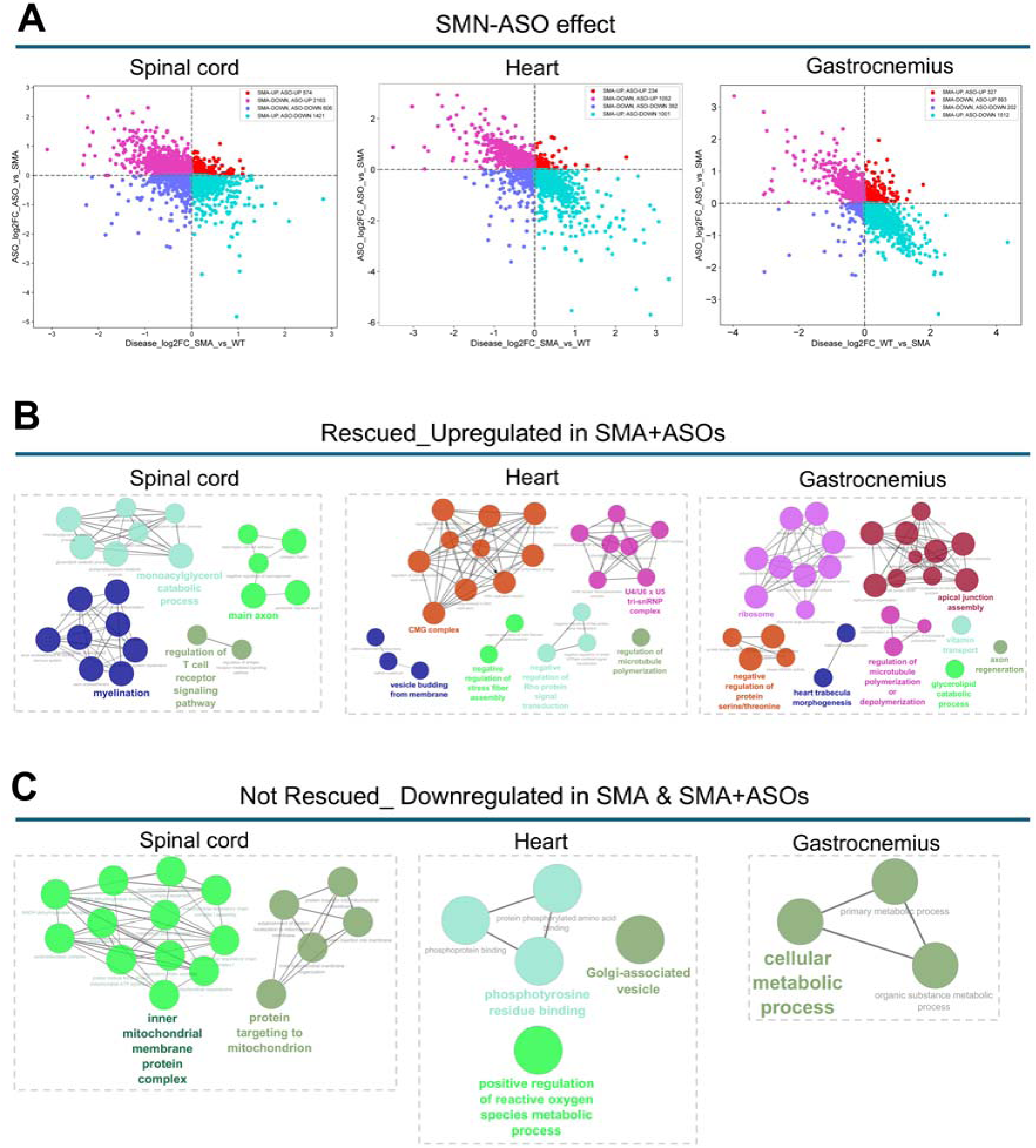
Functional characterization of rescued and persistent proteomic responses following SMN-ASO treatment. (A) Directional response classification plots showing protein abundance changes across spinal cord, heart, and gastrocnemius. Proteins are categorized based on combined WT versus SMA and SMA versus SMA+ASO comparisons into rescued (shift toward WT), persistent (remaining altered without directional reversal), and no change in ASO. (B) STRING network analysis of proteins rescued from SMA-associated upregulation following SMN-ASO treatment in spinal cord, heart, and gastrocnemius, visualized in Cytoscape with ClueGO functional grouping. Enriched pathways include immune-related processes in spinal cord, vesicle-mediated transport and cytoskeletal organization in heart, and synaptic, neuromuscular junction, and metabolic pathways in gastrocnemius. (C) STRING network analysis of proteins persistently downregulated in both SMA and SMA+ASO conditions, highlighting resistant pathways across tissues. Networks reveal enrichment of mitochondrial and metabolic processes, including inner mitochondrial membrane organization in spinal cord, vesicle-associated and redox-related pathways in heart, and cellular metabolic processes in gastrocnemius. Full list of rescued, persistent, no change in ASO and ASO-responsive-only targets per organ is provided in Supplementary Tables S7-S9.

Rescue-associated pathways differed across tissues, reflecting organ-specific responses to SMN restoration (Figure 5A-B; Supplementary Figure S6A; Supplementary Tables S6-S9). In spinal cord (Figure 5B), rescued upregulated proteins were enriched for immune-related pathways. In heart (Figure 5B), rescued categories included vesicle-mediated transport and cytoskeletal organization. In gastrocnemius (Figure 5B), rescued proteins were enriched for synaptic and neuromuscular junction-related pathways as well as metabolic processes.

A substantial subset of SMA-associated proteins remained uncorrected following SMN-ASO treatment (Figure 5A, C; Supplementary Figure S6B; Supplementary Tables S6-S9). Persistent downregulated proteins were enriched in mitochondrial respiratory chain and oxidative phosphorylation pathways in spinal cord and heart (Figure 5A, C; Supplementary Figure S6B). Persistent upregulated proteins were associated with stress-related and proteostasis pathways (Supplementary Figure S6B; Supplementary Tables S6-S9).

These findings indicate that SMN-ASO treatment selectively restores specific functional pathways even within the subgroup of SMA+ASO showing the clearest proteomic shift toward the WT/HET state, while key mitochondrial and stress-associated processes remain resistant to molecular correction.

## Discussion

Partial SMN restoration reshaped the proteomic landscape of SMA tissues in a tissue-dependent manner; however, this reorganization was selective rather than global. While subsets of inflammatory, synaptic, and structural pathways showed directional normalization, a substantial fraction of mitochondrial and metabolic alterations remained unresolved. These findings indicate that SMN-ASO treatment induces partial proteome reconfiguration rather than complete molecular restoration.

This incomplete reversibility was most evident in heart and spinal cord, where proteins associated with oxidative phosphorylation and mitochondrial organization remained persistently dysregulated. Previous transcriptomic studies in SMA spinal cord have consistently identified RNA-processing and splicing defects as primary consequences of SMN deficiency [54, 55], together with alterations in synaptic development and tissue organization [56, 57]. More recent multi-omics and single-cell analyses have expanded this view to include impaired protein synthesis, metabolic dysfunction, and vascular-associated signatures [58, 59]. Our data extend these observations to the protein level at the symptomatic stage, revealing coordinated mitochondrial and ER/Golgi-associated modules alongside axonogenesis- and glial differentiation-related networks.

Importantly, metabolic and mitochondrial signatures identified in spinal cord were paralleled in heart and gastrocnemius, supporting a multi-organ dimension of SMA pathology. Proteomic network analyses have previously implicated lipid metabolism and β-oxidation pathways in SMA [60], and early transcriptomic studies have suggested broader oxidoreductase and metabolic perturbations across tissues [61]. The present atlas integrates these findings across central and peripheral compartments, highlighting convergent metabolic vulnerability despite tissue-specific proteome remodeling.

The persistence of mitochondrial pathways following partial SMN restoration may indicate that secondary metabolic adaptations may not be fully reversible once established, particularly at symptomatic stages. This is consistent with biochemical observations showing incomplete normalization of redox-regulatory systems despite partial reduction of oxidative damage [24], as well as with evidence from other tissues where mitochondrial regulatory programs remain uncoupled from SMN partial recovery [25]. However, this interpretation should also be considered in light of the treatment paradigm used here in which SMN-ASO was administered systemically at a suboptimal dose designed to achieve partial rather than complete restoration of SMN levels [24, 33]. Thus, the incomplete proteomic rescue observed across tissues likely reflects both limited molecular correction and tissue-specific differences in the capacity to recover from established SMN deficiency.

An additional limitation of the rescue analysis is that downstream classification of rescued and persistent proteins was performed on the SMA+ASO samples showing the clearest proteomic shift toward the WT/HET state within each tissue. This stratified approach was chosen to assess treatment-associated molecular normalization in the context of heterogeneous responses within the SMA+ASO cohort, but it does not capture the full spectrum of proteomic behavior across all treated SMA animals. Accordingly, the rescue patterns described here should be interpreted as reflecting a subgroup with greater proteomic normalization rather than the entire SMA+ASO cohort. Nevertheless, because the analysis was performed at the level of whole-organ unbiased proteomics, the coordinated repositioning of multiple proteins and pathways across independent tissues supports the biological relevance of the observed rescue signatures despite the limited number of SMA+ASO subgroup that shifted towards WT/HET proteome distribution.

Overall, the heterogeneous degree of molecular rescue observed across tissues is compatible with the clinical variability in response to SMN-targeted therapies [19, 20, 62, 63]. Differences in treatment timing, tissue-specific vulnerability, and the extent of pre-existing remodeling are likely to influence the reversibility of molecular phenotypes [23].

## Conclusions

These data define a systemic yet heterogeneous proteomic response to SMN deficiency across neuromuscular organs, with peripheral tissues exhibiting broader molecular alterations than spinal cord at the symptomatic stage. Although SMN-ASO treatment partially repositioned tissue proteomes, mitochondrial and metabolic pathways remained incompletely normalized. In the context of previous biochemical and transcriptomic studies, these findings reinforce the view that mitochondrial and redox-associated dysfunction represent recurring components of SMA pathology and further support future studies exploring combinatorial therapeutic strategies.

## Supporting information

Volcano matrices Spinal cord

Volcano matrices Heart

Volcano matrices Gastrocnemius

WTvsSMA Intersection SC HR GC

HETvsSMA Intersection SC HR GC

ASO rescue summary overlap per organ

Rescue effect Spinal cord

Rescue effect Heart

Rescue effect Gastrocnemius

## Data Availability

Raw mass spectrometry data are available by the authors upon reasonable request. Processed outputs are included in the Supplementary material accompanying this preprint. Detailed experimental protocols are available in our previous works [24–26].

## Author contributions

**SV:** involved in conceptualization, performed all experiments, performed the proteomics workflow, performed statistical and bioinformatic analyses, wrote initial draft, generated the figures. **SM:** performed proteomics workflow, performed initial statistical analyses, involved in writing the methods, contributed to resources and funding. **BW:** involved in conceptualization, supervised the work, reviewed and edited the draft, contributed to funding acquisition.

## Acknowledgements

The work has been funded by the European Union’s Horizon 2020 Marie Skłodowska-Curie Program (project 956185; SMABEYOND) and the Center for Molecular Medicine Cologne (project C18) to BW and supported by the large instrument grant INST 216/1163-1 FUGG by the German Research Foundation (DFG Großgeräteantrag), to SM. We thank IONIS Pharmaceuticals for providing the SMN-ASOs and Roman Rombo for technical assistance in animal husbandry and treating the mice. The graphical abstract created using BioRender.com.

## References

[1] Prior T, Leach M, Finanger E. Spinal Muscular Atrophy. 2000 Feb 24 [updated 2024 Sep 19]. In: GeneReviews® [Internet]. Seattle (WA): University of Washington, Seattle. Available from: https://www.ncbi.nlm.nih.gov/books/NBK1352/.

[2] Lefebvre S, Bürglen L, Reboullet S, Clermont O, Burlet P, Viollet L, et al., Identification and characterization of a spinal muscular atrophy-determining gene, Cell 80 (1995) 155–65, doi:10.1016/0092-8674(95)90460-3.

[3] Lorson CL, Hahnen E, Androphy EJ, Wirth B, A single nucleotide in the SMN gene regulates splicing and is responsible for spinal muscular atrophy, Proc Natl Acad Sci U S A 96 (1999) 6307–11, doi:10.1073/pnas.96.11.6307.

[4] Singh RN, Howell MD, Ottesen EW, Singh NN, Diverse role of survival motor neuron protein, Biochim Biophys Acta Gene Regul Mech 1860 (2017) 299–315, doi:10.1016/j.bbagrm.2016.12.008.

[5] Yeo CJJ, Darras BT, Overturning the Paradigm of Spinal Muscular Atrophy as Just a Motor Neuron Disease, Pediatr Neurol 109 (2020) 12–9, doi:10.1016/j.pediatrneurol.2020.01.003.

[6] Hamilton G, Gillingwater TH, Spinal muscular atrophy: going beyond the motor neuron, Trends Mol Med 19 (2013) 40–50, doi:10.1016/j.molmed.2012.11.002.

[7] Shababi M, Habibi J, Yang HT, Vale SM, Sewell WA, Lorson CL, Cardiac defects contribute to the pathology of spinal muscular atrophy models, Human Molecular Genetics 19 (2010) 4059–71, doi:10.1093/hmg/ddq329.

[8] Ripolone M, Ronchi D, Violano R, Vallejo D, Fagiolari G, Barca E, et al., Impaired Muscle Mitochondrial Biogenesis and Myogenesis in Spinal Muscular Atrophy, JAMA Neurol 72 (2015) 666–75, doi:10.1001/jamaneurol.2015.0178.

[9] Leow DM-K, Ng YK, Wang LC, Koh HWL, Zhao T, Khong ZJ, et al., Hepatocyte-intrinsic SMN deficiency drives metabolic dysfunction and liver steatosis in spinal muscular atrophy, The Journal of Clinical Investigation 134 (2024) doi:10.1172/JCI173702.

[10] Hua Y, Sahashi K, Rigo F, Hung G, Horev G, Bennett CF, et al., Peripheral SMN restoration is essential for long-term rescue of a severe spinal muscular atrophy mouse model, Nature 478 (2011) 123–6, doi:10.1038/nature10485.

[11] Deguise MO, Baranello G, Mastella C, Beauvais A, Michaud J, Leone A, et al., Abnormal fatty acid metabolism is a core component of spinal muscular atrophy, Ann Clin Transl Neurol 6 (2019) 1519–32, doi:10.1002/acn3.50855.

[12] Boyer JG, Deguise MO, Murray LM, Yazdani A, De Repentigny Y, Boudreau-Larivière C, et al., Myogenic program dysregulation is contributory to disease pathogenesis in spinal muscular atrophy, Hum Mol Genet 23 (2014) 4249–59, doi:10.1093/hmg/ddu142.

[13] Ling KK, Gibbs RM, Feng Z, Ko CP, Severe neuromuscular denervation of clinically relevant muscles in a mouse model of spinal muscular atrophy, Hum Mol Genet 21 (2012) 185–95, doi:10.1093/hmg/ddr453.

[14] Shababi M, Habibi J, Yang HT, Vale SM, Sewell WA, Lorson CL, Cardiac defects contribute to the pathology of spinal muscular atrophy models, Hum Mol Genet 19 (2010) 4059–71, doi:10.1093/hmg/ddq329.

[15] Chemello F, Pozzobon M, Tsansizi LI, Varanita T, Quintana-Cabrera R, Bonesso D, et al., Dysfunctional mitochondria accumulate in a skeletal muscle knockout model of Smn1, the causal gene of spinal muscular atrophy, Cell Death & Disease 14 (2023) 162, doi:10.1038/s41419-023-05573-x.

[16] Acsadi G, Lee I, Li X, Khaidakov M, Pecinova A, Parker GC, et al., Mitochondrial dysfunction in a neural cell model of spinal muscular atrophy, J Neurosci Res 87 (2009) 2748–56, doi:10.1002/jnr.22106.

[17] Zilio E, Piano V, Wirth B, Mitochondrial Dysfunction in Spinal Muscular Atrophy, International Journal of Molecular Sciences 23 (2022) 10878, doi:10.3390/ijms231810878.

[18] James R, Chaytow H, Ledahawsky LM, Gillingwater TH, Revisiting the role of mitochondria in spinal muscular atrophy, Cell Mol Life Sci 78 (2021) 4785–804, doi:10.1007/s00018-021-03819-5.

[19] Mercuri E, Darras BT, Chiriboga CA, Day JW, Campbell C, Connolly AM, et al., Nusinersen versus Sham Control in Later-Onset Spinal Muscular Atrophy, N Engl J Med 378 (2018) 625–35, doi:10.1056/NEJMoa1710504.

[20] Mendell JR, Al-Zaidy S, Shell R, Arnold WD, Rodino-Klapac LR, Prior TW, et al., Single-Dose Gene-Replacement Therapy for Spinal Muscular Atrophy, N Engl J Med 377 (2017) 1713–22, doi:10.1056/NEJMoa1706198.

[21] Passini MA, Bu J, Richards AM, Kinnecom C, Sardi SP, Stanek LM, et al., Antisense oligonucleotides delivered to the mouse CNS ameliorate symptoms of severe spinal muscular atrophy, Sci Transl Med 3 (2011) 72ra18, doi:10.1126/scitranslmed.3001777.

[22] Porensky PN, Mitrpant C, McGovern VL, Bevan AK, Foust KD, Kaspar BK, et al., A single administration of morpholino antisense oligomer rescues spinal muscular atrophy in mouse, Hum Mol Genet 21 (2012) 1625–38, doi:10.1093/hmg/ddr600.

[23] Wirth B, Spinal Muscular Atrophy: In the Challenge Lies a Solution, Trends Neurosci 44 (2021) 306–22, doi:10.1016/j.tins.2020.11.009.

[24] Vrettou S, Wirth B, Organ-specific redox imbalances in spinal muscular atrophy mice are partially rescued by SMN antisense oligonucleotides, FEBS Letters (2026) doi:10.1002/1873-3468.70303.

[25] Vrettou S, Müller S, Wirth B, SMN deficiency disrupts hepatic mitochondrial iron homeostasis and NRF2-dependent redox control in spinal muscular atrophy, bioRxiv (2026) 2026.01.08.698518, doi:10.64898/2026.01.08.698518.

[26] Vrettou S, Müller S, Wirth B, Proteomics dataset of liver tissue from spinal muscular atrophy, heterozygous, and wild-type mice, enabling pathway identification, Data in Brief (2026) 112632, doi:10.1016/j.dib.2026.112632.

[27] Riessland M, Ackermann B, Förster A, Jakubik M, Hauke J, Garbes L, et al., SAHA ameliorates the SMA phenotype in two mouse models for spinal muscular atrophy, Hum Mol Genet 19 (2010) 1492–506, doi:10.1093/hmg/ddq023.

[28] Muinos-Bühl A, Rombo R, Janzen E, Ling KK, Hupperich K, Rigo F, et al., Combinatorial ASO-mediated therapy with low dose SMN and the protective modifier Chp1 is not sufficient to ameliorate SMA pathology hallmarks, Neurobiol Dis 171 (2022) 105795, doi:10.1016/j.nbd.2022.105795.

[29] Hsieh-Li HM, Chang JG, Jong YJ, Wu MH, Wang NM, Tsai CH, et al., A mouse model for spinal muscular atrophy, Nat Genet 24 (2000) 66–70, doi:10.1038/71709.

[30] Torres-Benito L, Schneider S, Rombo R, Ling KK, Grysko V, Upadhyay A, et al., NCALD Antisense Oligonucleotide Therapy in Addition to Nusinersen further Ameliorates Spinal Muscular Atrophy in Mice, Am J Hum Genet 105 (2019) 221–30, doi:10.1016/j.ajhg.2019.05.008.

[31] Muiños-Bühl A, Rombo R, Ling KK, Zilio E, Rigo F, Bennett CF, et al., Long-Term SMN- and Ncald-ASO Combinatorial Therapy in SMA Mice and NCALD-ASO Treatment in hiPSC-Derived Motor Neurons Show Protective Effects, Int J Mol Sci 24 (2023) doi:10.3390/ijms24044198.

[32] Hua Y, Vickers TA, Okunola HL, Bennett CF, Krainer AR, Antisense masking of an hnRNP A1/A2 intronic splicing silencer corrects SMN2 splicing in transgenic mice, Am J Hum Genet 82 (2008) 834–48, doi:10.1016/j.ajhg.2008.01.014.

[33] Hosseinibarkooie S, Peters M, Torres-Benito L, Rastetter RH, Hupperich K, Hoffmann A, et al., The Power of Human Protective Modifiers: PLS3 and CORO1C Unravel Impaired Endocytosis in Spinal Muscular Atrophy and Rescue SMA Phenotype, Am J Hum Genet 99 (2016) 647–65, doi:10.1016/j.ajhg.2016.07.014.

[34] Demichev V, Messner CB, Vernardis SI, Lilley KS, Ralser M, DIA-NN: neural networks and interference correction enable deep proteome coverage in high throughput, Nat Methods 17 (2020) 41–4, doi:10.1038/s41592-019-0638-x.

[35] Tyanova S, Temu T, Sinitcyn P, Carlson A, Hein MY, Geiger T, et al., The Perseus computational platform for comprehensive analysis of (prote)omics data, Nature Methods 13 (2016) 731–40, doi:10.1038/nmeth.3901.

[36] Nolte H, MacVicar TD, Tellkamp F, Krüger M, Instant Clue: A Software Suite for Interactive Data Visualization and Analysis, Scientific Reports 8 (2018) 12648, doi:10.1038/s41598-018-31154-6.

[37] Tang D, Chen M, Huang X, Zhang G, Zeng L, Zhang G, et al., SRplot: A free online platform for data visualization and graphing, PLOS ONE 18 (2023) e0294236, doi:10.1371/journal.pone.0294236.

[38] Pellizzoni L, Kataoka N, Charroux B, Dreyfuss G, A novel function for SMN, the spinal muscular atrophy disease gene product, in pre-mRNA splicing, Cell 95 (1998) 615–24, doi:10.1016/s0092-8674(00)81632-3.

[39] Lefebvre S, Bürglen L, Frézal J, Munnich A, Melki J, The Role of the SMN Gene in Proximal Spinal Muscular Atrophy, Human Molecular Genetics 7 (1998) 1531–6, doi:10.1093/hmg/7.10.1531.

[40] Jablonka S, Rossoll W, Schrank B, Sendtner M, The role of SMN in spinal muscular atrophy, Journal of Neurology 247 (2000) I37–I42, doi:10.1007/s004150050555.

[41] Torres-Benito L, Neher MF, Cano R, Ruiz R, Tabares L, SMN Requirement for Synaptic Vesicle, Active Zone and Microtubule Postnatal Organization in Motor Nerve Terminals, PLOS ONE 6 (2011) e26164, doi:10.1371/journal.pone.0026164.

[42] Oprea GE, Kröber S, McWhorter ML, Rossoll W, Müller S, Krawczak M, et al., Plastin 3 is a protective modifier of autosomal recessive spinal muscular atrophy, Science 320 (2008) 524–7, doi:10.1126/science.1155085.

[43] Nölle A, Zeug A, van Bergeijk J, Tönges L, Gerhard R, Brinkmann H, et al., The spinal muscular atrophy disease protein SMN is linked to the rho-kinase pathway via profilin, Human Molecular Genetics 20 (2011) 4865–78, doi:10.1093/hmg/ddr425.

[44] Hensel N, Claus P, The Actin Cytoskeleton in SMA and ALS: How Does It Contribute to Motoneuron Degeneration?, Neuroscientist 24 (2018) 54–72, doi:10.1177/1073858417705059.

[45] Bowerman M, Shafey D, Kothary R, Smn Depletion Alters Profilin II Expression and Leads to Upregulation of the RhoA/ROCK Pathway and Defects in Neuronal Integrity, Journal of Molecular Neuroscience 32 (2007) 120–31, doi:10.1007/s12031-007-0024-5.

[46] Walsh MB, Janzen E, Wingrove E, Hosseinibarkooie S, Muela NR, Davidow L, et al., Genetic modifiers ameliorate endocytic and neuromuscular defects in a model of spinal muscular atrophy, BMC Biol 18 (2020) 127, doi:10.1186/s12915-020-00845-w.

[47] Riessland M, Kaczmarek A, Schneider S, Swoboda KJ, Löhr H, Bradler C, et al., Neurocalcin Delta Suppression Protects against Spinal Muscular Atrophy in Humans and across Species by Restoring Impaired Endocytosis, Am J Hum Genet 100 (2017) 297–315, doi:10.1016/j.ajhg.2017.01.005.

[48] Janzen E, Mendoza-Ferreira N, Hosseinibarkooie S, Schneider S, Hupperich K, Tschanz T, et al., CHP1 reduction ameliorates spinal muscular atrophy pathology by restoring calcineurin activity and endocytosis, Brain 141 (2018) 2343–61, doi:10.1093/brain/awy167.

[49] Dimitriadi M, Derdowski A, Kalloo G, Maginnis MS, O’Hern P, Bliska B, et al., Decreased function of survival motor neuron protein impairs endocytic pathways, Proc Natl Acad Sci U S A 113 (2016) E4377–86, doi:10.1073/pnas.1600015113.

[50] Wadman RI, Vrancken AFJE, van den Berg LH, van der Pol WL, Dysfunction of the neuromuscular junction in spinal muscular atrophy types 2 and 3, Neurology 79 (2012) 2050–5, doi:10.1212/WNL.0b013e3182749eca.

[51] Murray LM, Comley LH, Thomson D, Parkinson N, Talbot K, Gillingwater TH, Selective vulnerability of motor neurons and dissociation of pre- and post-synaptic pathology at the neuromuscular junction in mouse models of spinal muscular atrophy, Human Molecular Genetics 17 (2008) 949–62, doi:10.1093/hmg/ddm367.

[52] Courtney NL, Mole AJ, Thomson AK, Murray LM, Reduced P53 levels ameliorate neuromuscular junction loss without affecting motor neuron pathology in a mouse model of spinal muscular atrophy, Cell Death & Disease 10 (2019) 515, doi:10.1038/s41419-019-1727-6.

[53] Arnold WD, Severyn S, Zhao S, Kline D, Linsenmayer M, Kelly K, et al., Persistent neuromuscular junction transmission defects in adults with spinal muscular atrophy treated with nusinersen, BMJ Neurol Open 3 (2021) e000164, doi:10.1136/bmjno-2021-000164.

[54] Jangi M, Fleet C, Cullen P, Gupta SV, Mekhoubad S, Chiao E, et al., SMN deficiency in severe models of spinal muscular atrophy causes widespread intron retention and DNA damage, Proceedings of the National Academy of Sciences 114 (2017) E2347–E56, doi:10.1073/pnas.1613181114.

[55] Doktor TK, Hua Y, Andersen HS, Brøner S, Liu YH, Wieckowska A, et al., RNA-sequencing of a mouse-model of spinal muscular atrophy reveals tissue-wide changes in splicing of U12-dependent introns, Nucleic Acids Res 45 (2017) 395–416, doi:10.1093/nar/gkw731.

[56] Zhang Z, Pinto AM, Wan L, Wang W, Berg MG, Oliva I, et al., Dysregulation of synaptogenesis genes antecedes motor neuron pathology in spinal muscular atrophy, Proceedings of the National Academy of Sciences 110 (2013) 19348–53, doi:10.1073/pnas.1319280110.

[57] Murray LM, Lee S, Bäumer D, Parson SH, Talbot K, Gillingwater TH, Pre-symptomatic development of lower motor neuron connectivity in a mouse model of severe spinal muscular atrophy, Hum Mol Genet 19 (2010) 420–33, doi:10.1093/hmg/ddp506.

[58] Sun J, Qiu J, Yang Q, Ju Q, Qu R, Wang X, et al., Single-cell RNA sequencing reveals dysregulation of spinal cord cell types in a severe spinal muscular atrophy mouse model, PLOS Genetics 18 (2022) e1010392, doi:10.1371/journal.pgen.1010392.

[59] Nichterwitz S, Nijssen J, Storvall H, Schweingruber C, Comley LH, Allodi I, et al., LCM-seq reveals unique transcriptional adaptation mechanisms of resistant neurons and identifies protective pathways in spinal muscular atrophy, Genome Res 30 (2020) 1083–96, doi:10.1101/gr.265017.120.

[60] Tapken I, Schweitzer T, Paganin M, Schüning T, Detering NT, Sharma G, et al., The systemic complexity of a monogenic disease: the molecular network of spinal muscular atrophy, Brain 148 (2025) 580–96, doi:10.1093/brain/awae272.

[61] Wu L, Sun J, Wang L, Chen Z, Guan Z, Du L, et al., Whole-transcriptome sequencing in neural and non-neural tissues of a mouse model identifies miR-34a as a key regulator in SMA pathogenesis, Mol Ther Nucleic Acids 36 (2025) 102490, doi:10.1016/j.omtn.2025.102490.

[62] Pane M, Coratti G, Sansone VA, Messina S, Bruno C, Catteruccia M, et al., Nusinersen in type 1 spinal muscular atrophy: Twelve-month real-world data, Annals of Neurology 86 (2019) 443–51, doi:10.1002/ana.25533.

[63] Darras BT, Chiriboga CA, Iannaccone ST, Swoboda KJ, Montes J, Mignon L, et al., Nusinersen in later-onset spinal muscular atrophy: Long-term results from the phase 1/2 studies, Neurology 92 (2019) e2492–e506, doi:10.1212/wnl.0000000000007527.

