## Supplementary Figures S1-S6 for "System-Wide Proteomic Remodeling in Spinal Muscular Atrophy Reveals Tissue-Specific Responses and Partial Rescue by SMN Restoration"

### Content

- **Supplementary Figure S1. Differential proteome analysis across genotypes and cross-tissue overlap.**
- **Supplementary Figure S2. Cross-tissue convergence of SMA-associated proteome alterations in heart and gastrocnemius.**
- **Supplementary Figure S3. Network organization of upregulated proteomes in WT versus SMA comparisons across tissues.**
- **Supplementary Figure S4. Network organization of upregulated proteomes in HET versus SMA comparisons across tissues.**
- **Supplementary Figure S5. Proteomic profiling of heterozygous (HET) mice following SMN-ASO treatment across neuromuscular tissues.**
- **Supplementary Figure S6. Functional characterization of rescue-associated and persistent proteomic alterations following SMN-ASO treatment.**

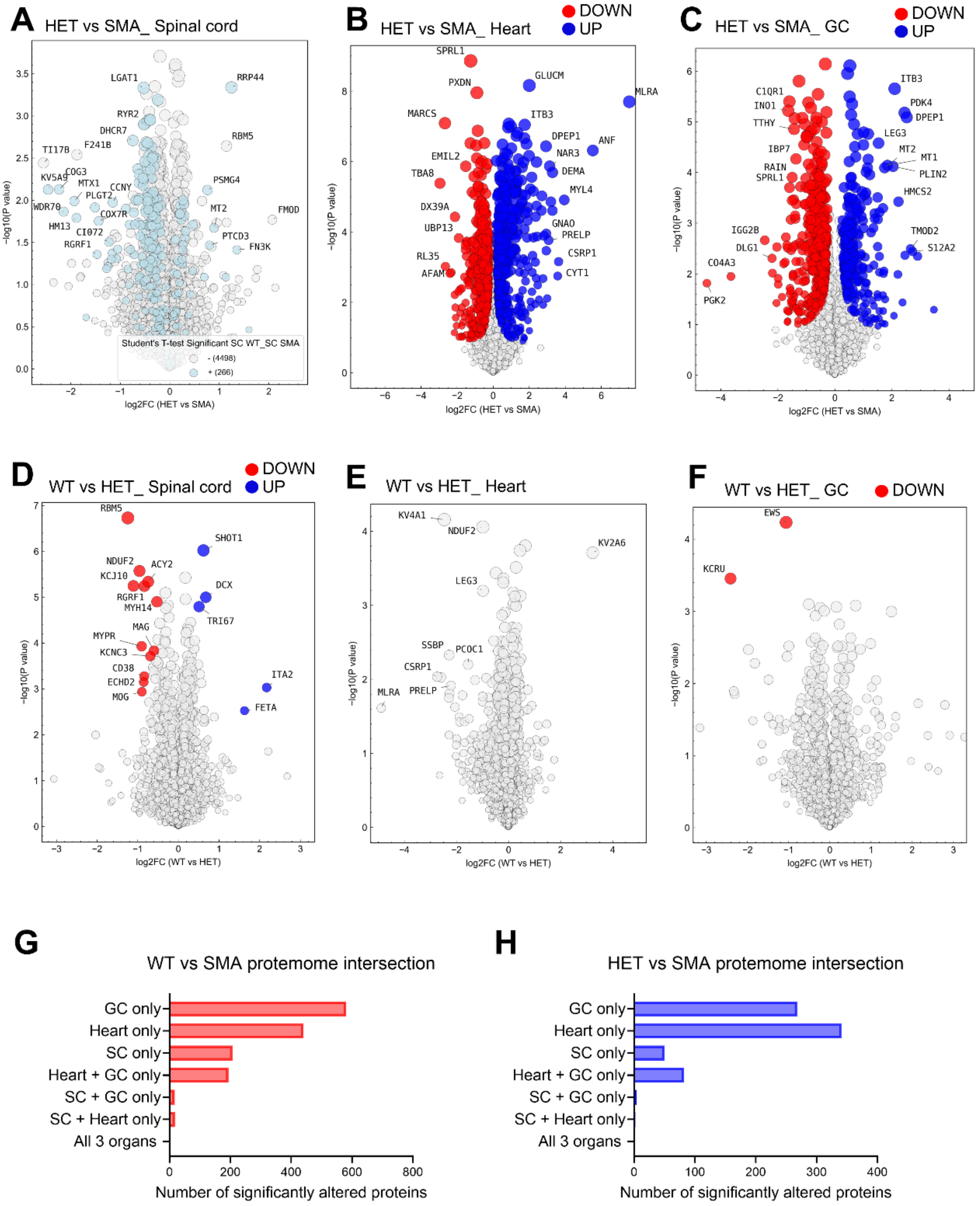


**Supplementary Figure S1. Differential proteome analysis across genotypes and cross-tissue overlap.** (A-C) Volcano plots of differential protein abundance for HET vs SMA in spinal cord (A), heart (B), and gastrocnemius muscle (C), generated in Perseus. Proteins passing significance thresholds in the HET vs SMA comparison are shown as significantly downregulated (red) or upregulated (blue). In addition, proteins significantly altered in the corresponding WT vs SMA comparison but not reaching significance in HET vs SMA are overlaid to visualize shared directional trends between comparisons. No proteins passed the applied thresholds in spinal cord, whereas heart and gastrocnemius showed substantial differential proteome remodeling. (D-F) Volcano plots of WT vs HET comparisons in spinal cord (D), heart (E), and gastrocnemius (F), showing limited proteome alterations between these genotypes. (G, H) Intersection analysis of significantly altered proteins across tissues for WT vs SMA (G) and HET vs SMA (H), illustrating that most proteome changes are tissue-specific, with limited overlap between organs. These analyses show that HET vs SMA shares directional proteomic features with WT vs SMA, particularly in heart and gastrocnemius, despite fewer proteins passing statistical thresholds in the HET comparison.

A

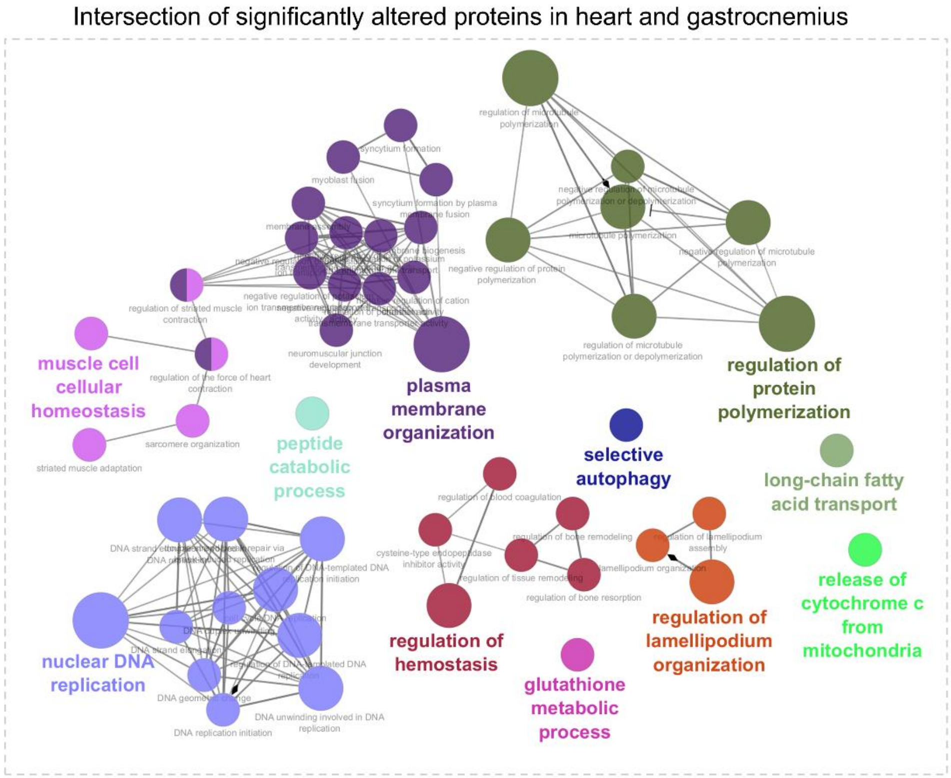


**Supplementary Figure S2. Cross-tissue convergence of SMA-associated proteome alterations in heart and gastrocnemius.** (A) STRING network analysis of proteins significantly altered in both heart and gastrocnemius (WT vs SMA comparison), visualized in Cytoscape with ClueGO functional grouping. Shared proteins were identified based on UniProt ID overlap between tissues. Network clustering reveals convergent biological processes, including plasma membrane organization, muscle cell homeostasis, DNA replication, protein polymerization, autophagy, mitochondrial function, and metabolic pathways. These data indicate pathway-level convergence between peripheral tissues despite limited overlap at the individual protein level.

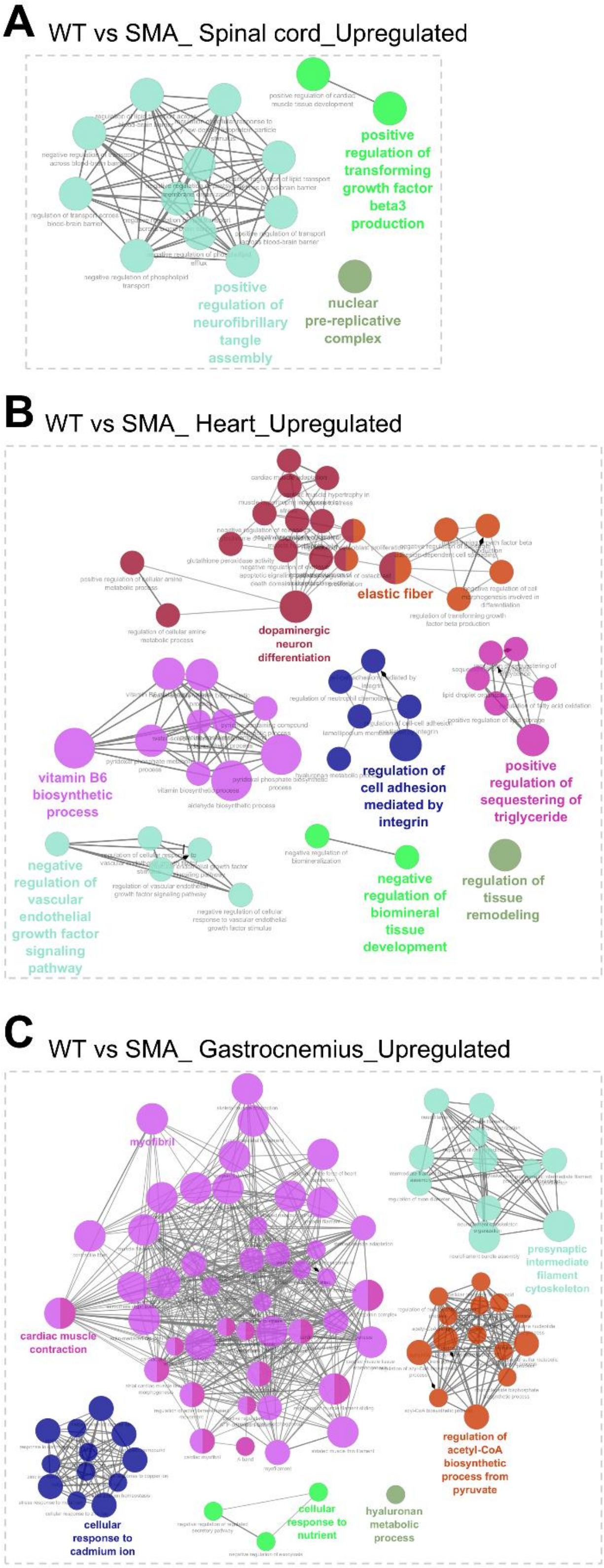


**Supplementary Figure S3. Network organization of upregulated proteomes in WT versus SMA comparisons across tissues.** (A-C) STRING network analysis of proteins significantly upregulated in WT vs SMA comparisons in spinal cord (A), heart (B), and gastrocnemius muscle (C), visualized in Cytoscape with ClueGO functional grouping. Networks reveal tissue-specific enrichment of biological processes, including growth factor–related signaling and nuclear processes in spinal cord (A); extracellular matrix organization, lipid metabolism, tissue remodeling, and signaling pathways in heart (B); and muscle contraction, cytoskeletal organization, metabolic processes, and stress responses in gastrocnemius (C). These analyses complement downregulated network findings by highlighting distinct, tissue-specific upregulated pathways associated with SMN deficiency.

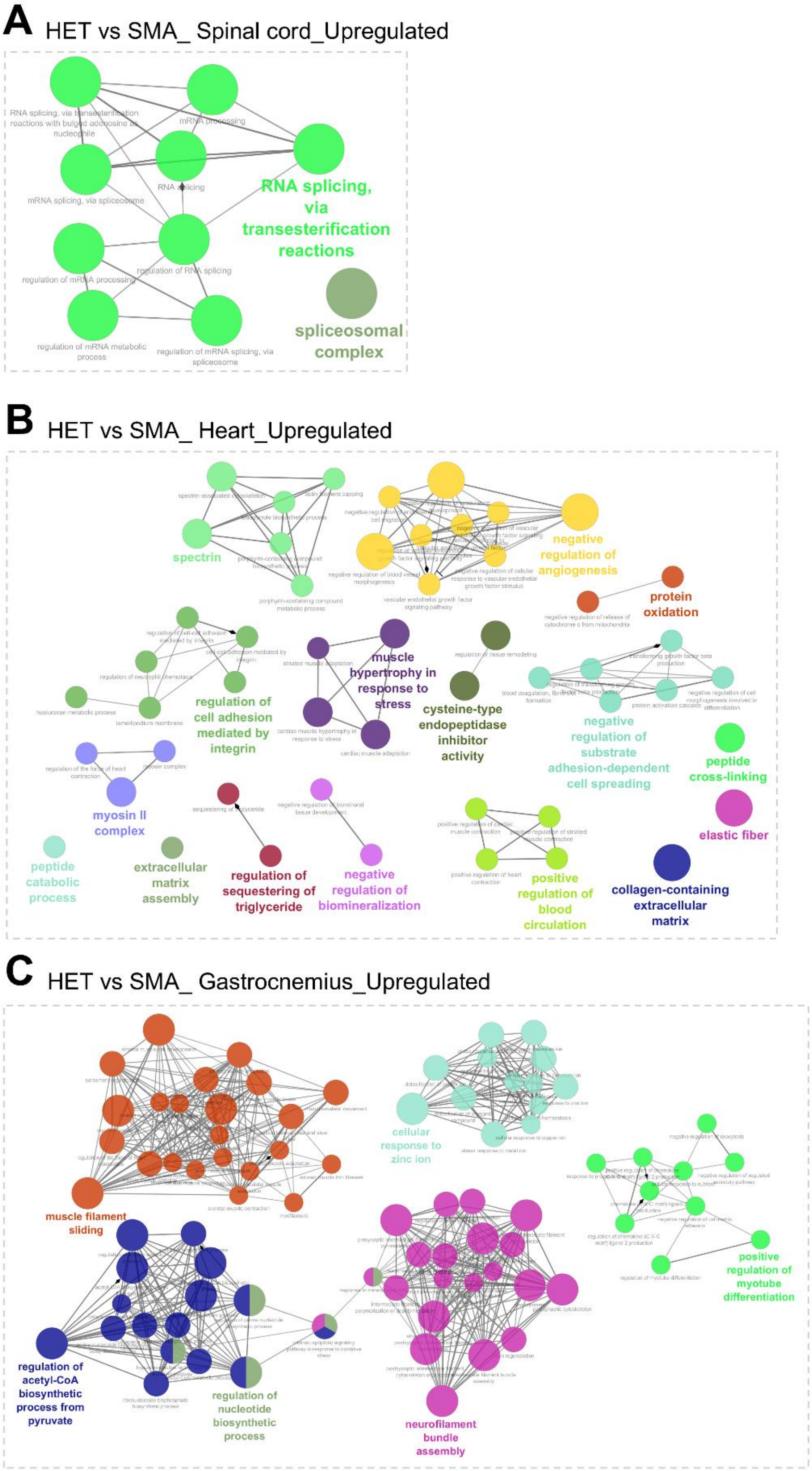


**Supplementary Figure S4. Network organization of upregulated proteomes in HET versus SMA comparisons across tissues.** (A-C) STRING network analysis of proteins significantly upregulated in HET vs SMA comparisons in spinal cord (A), heart (B), and gastrocnemius muscle (C), visualized in Cytoscape with ClueGO functional grouping. Networks reveal tissue-specific enrichment of biological processes, including RNA splicing and spliceosomal components in spinal cord (A); extracellular matrix organization, cell adhesion, angiogenesis, and metabolic processes in heart (B); and muscle contraction, cytoskeletal organization, metabolic pathways, and differentiation processes in gastrocnemius (C). These analyses indicate that SMN dosage-dependent proteome changes involve distinct functional programs across tissues, with partial overlap in RNA processing, structural, and metabolic pathways.

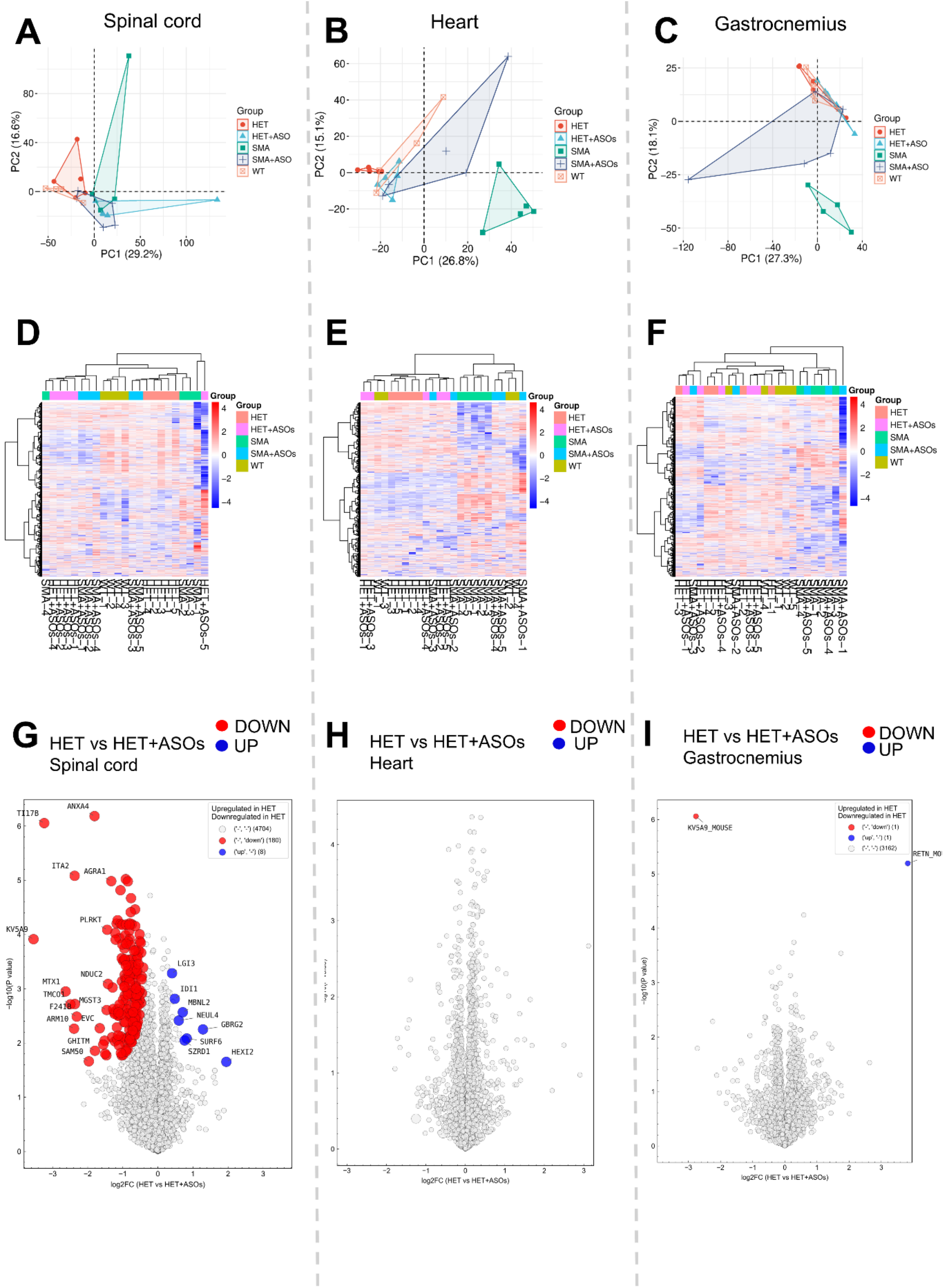


**Supplementary Figure S5. Proteomic profiling of heterozygous (HET) mice following SMN-ASO treatment across neuromuscular tissues.** (A-C) Principal component analysis (PCA) of spinal cord (A), heart (B), and gastrocnemius muscle (C) including all experimental groups (WT, HET, SMA, HET+ASOs, SMA+ASOs). HET and HET+ASO samples cluster closely across tissues, indicating minimal global proteome shifts upon ASO treatment in heterozygous animals. (D-F) Unsupervised hierarchical clustering of protein abundance across spinal cord (D), heart (E), and gastrocnemius (F) for all groups. Samples primarily cluster according to genotype, with HET and HET+ASO groups displaying similar proteomic profiles. (G-I) Volcano plots showing differential protein abundance between HET and HET+ASO groups in spinal cord (G), heart (H), and gastrocnemius (I). Significantly downregulated (red) and upregulated (blue) proteins are indicated based on permutation-based FDR thresholds in Perseus. Spinal cord exhibits a subset of significantly altered proteins, whereas heart and gastrocnemius show minimal to no changes following ASO treatment.

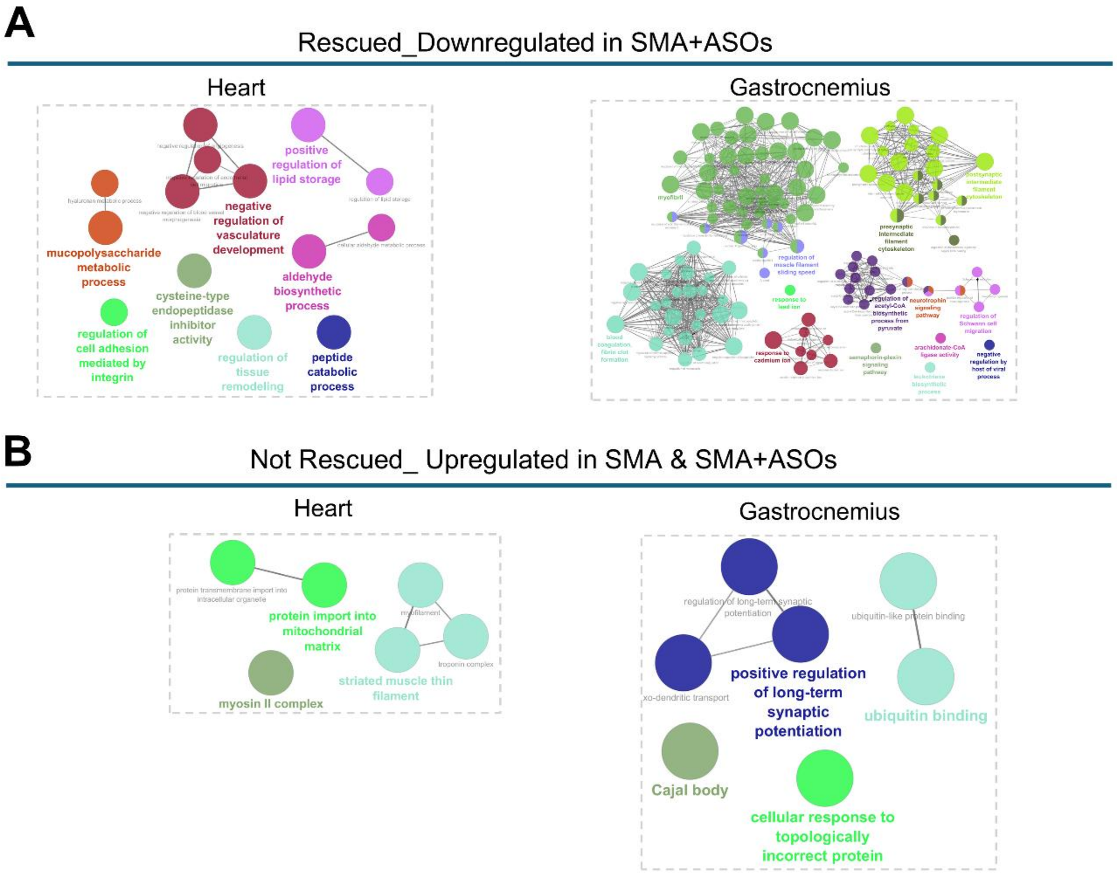


**Supplementary Figure S6. Functional characterization of rescue-associated and persistent proteomic alterations following SMN-ASO treatment.** (A) STRING network analysis of proteins that were significantly dysregulated in SMA and directionally reversed following SMN-ASO treatment, shown for heart (left) and gastrocnemius muscle (right). These rescue-associated networks represent proteins whose abundance shifted toward control levels after treatment. In spinal cord, rescue-associated proteins did not form significantly enriched or connected functional clusters. (B) STRING network analysis of proteins that remained dysregulated despite SMN-ASO treatment, shown for heart (left) and gastrocnemius muscle (right). These persistent networks represent pathways resistant to molecular correction following SMN restoration. In spinal cord, persistent proteins did not form associated functional networks.
